# Increased 20-HETE signaling suppresses neurovascular coupling after ischemic stroke in regions beyond the infarct

**DOI:** 10.1101/2021.08.25.457547

**Authors:** Zhenzhou Li, Heather L. McConnell, Teresa L. Stackhouse, Martin M. Pike, Wenri Zhang, Anusha Mishra

## Abstract

Neurovascular coupling, the process by which neuronal activity elicits increases in the local blood supply, is impaired in stroke patients in brain regions outside the infarct. Such impairment may contribute to neurological deterioration over time, but its mechanism is unknown. Using the middle cerebral artery occlusion (MCAO) model of stroke, we show that neuronal activity-evoked capillary dilation is reduced by ∼75% in the intact cortical tissue outside the infarct border. This decrease in capillary responsiveness was not explained by a decrease in local neuronal activity or a loss of vascular contractility. Inhibiting synthesis of the vasoconstrictive molecule 20-HETE, either by inhibiting its synthetic enzyme CYP450 ω-hydroxylases or by increasing nitric oxide (NO), which is a natural inhibitor of ω-hydroxylases, rescued activity-evoked capillary dilation. The capillary dilation unmasked by inhibiting 20-HETE was dependent on PGE2 activation of EP4 receptors, a vasodilatory pathway previously identified in healthy animals. Cortical 20-HETE levels were increased following MCAO, in agreement with data from stroke patients. Inhibition of ω-hydroxylases normalized 20-HETE levels *in vivo* and increased cerebral blood flow in the peri-infarct cortex. These data identify 20-HETE-dependent vasoconstriction as a mechanism underlying neurovascular coupling impairment after stroke. Our results suggest that the brain’s energy supply may be significantly reduced after stroke in regions previously believed to be asymptomatic and that ω-hydroxylase inhibition may restore healthy neurovascular coupling post-stroke.

## Introduction

Neuronal activity in the central nervous system is coupled to an increase in local blood flow [1, 2]. This process, known as neurovascular coupling, ensures that the increased energy demand of active neural tissue is met by a supply of metabolic substrates such as oxygen and glucose. Neurovascular coupling is impaired in many neurodegenerative conditions, including Alzheimer’s disease (AD), amyotrophic lateral sclerosis, and multiple sclerosis [3-6]. Impaired neurovascular coupling is also reported in patients diagnosed with stroke [7-9], especially strokes arising from occlusion of the middle cerebral artery [10]. Importantly, neurovascular coupling impairment is observed in stroke patients even when they are considered to be fully recovered, for up to several years following the stroke and in clinically asymptomatic regions outside the infarct lesion [8]. This impairment could result in energy deficits in otherwise healthy brain regions and thereby compromise neuronal health over time.

Ischemic injuries, including stroke, transient ischemic attacks, silent infarcts, and watershed microinfarcts, increase the risk of dementia by several fold [11-15], and this is not explained purely by common risk factors [16]. Conversely, patients with dementia often show signs of previous ischemic injuries in their brains [14, 15, 17]. Furthermore, cerebrovascular dysfunction is one of the earliest detectable changes in patients who develop cognitive dysfunction and dementia [18, 19]. Ischemia-induced impairment of neurovascular coupling is likely a major contributor to such cerebrovascular dysfunction, yet mechanisms underlying this impairment have not yet been studied.

In the healthy brain, neurovascular coupling at the capillary level occurs via an astrocyte-mediated pathway that relies on vasoactive metabolites of arachidonic acid. In response to neuronal activity, astrocytes synthesize arachidonic acid and metabolize it to form prostaglandin E2 (PGE2), which then acts on the endoperoxide 4 (EP4) receptor on contractile pericytes to induce capillary dilation [20]. Arachidonic acid can also be metabolized to the vasoconstrictive molecule 20-hydroxyeicosatetraenoic acid (20-HETE). The CYP450 ω-hydroxylase enzymes that synthesize 20-HETE are inhibited in physiologically healthy tissue, however, by activity-dependent production of nitric oxide (NO) [21, 22]. Thus, neuronal activity leads to a net dilation of capillaries via EP4 receptors.

Existing evidence suggests that increased levels of 20-HETE in the cerebrospinal fluid [23] and plasma [24, 25] of stroke patients predict a negative clinical prognosis. Further, an increase in 20-HETE synthesis occurs after experimental stroke in rodent models and contributes to a decrease in global cerebral blood flow (CBF) after stroke [26] and cortical spreading depression [27]. Evidence also suggests that neuronal and endothelial NO synthases are downregulated after ischemic vascular injury [28]. These findings led us to hypothesize that a decrease in NO disinhibits 20-HETE synthesis and thereby impairs neurovascular coupling after stroke.

Here, we applied transient middle cerebral artery occlusion (MCAO) and assayed neurovascular coupling in acute brain slices prepared one day after transient MCAO. We found that neuronal activity-induced capillary dilation is severely diminished in intact cortical tissue beyond the infarct border, a region we term the ‘peri-infarct,’ compared to the analogous region in the contralateral hemisphere or naïve brains. We show that impairment of neurovascular coupling is not due to decreased neuronal activity or vascular contractility, but rather due to increased synthesis of the vasoconstrictor 20-HETE. Importantly, we demonstrate that preventing 20-HETE synthesis can restore healthy neurovascular coupling *ex vivo* and elevate CBF *in vivo*, selectively in the peri-infarct cortex of the stroke hemisphere. These data suggest a mechanism for neurovascular coupling impairment after stroke in the intact cortex surrounding the infarct, mediated by 20-HETE synthesis.

## Materials and Methods

### Middle Cerebral Artery Occlusion (MCAO)

All experiments were approved by the Oregon Health & Science University Institutional Animal Care and Use Committee. Two-month-old Long-Evans rats (Charles River Laboratories) of either sex were randomly assigned to experimental (MCAO) or naïve groups. All animals were in healthy condition prior to manipulation. Rats were placed under isoflurane anesthesia (5% induction, 1.5% maintenance) in 30% oxygen-enriched air via mask. Body temperature was maintained at 37 ± 0.5°C throughout the procedure. Middle cerebral artery (MCA) occlusion was performed using a previously described method with slight modifications [29]. Briefly, a laser Doppler flowmeter (Moore Instruments) probe was affixed over the right parietal bone overlying the MCA territory to monitor changes in CBF. A ventral midline incision was made over the neck, the right common carotid artery (CCA) bifurcation was exposed by gentle dissection and tissue retraction, and the external carotid artery (ECA) was permanently ligated distal to the occipital artery using electrocautery, such that a short ECA stump remained attached to the bifurcation. The right CCA and internal carotid arteries (ICA) were temporarily closed with reversible slip knots before an arteriotomy was made in the ECA stump. A silicone-coated 5.0 nylon monofilament (Doccol Corporation) appropriate for the weight of the rat, as suggested by the manufacturer, was inserted into the ICA via the arteriotomy and gently advanced to the ICA/MCA bifurcation to occlude CBF to the MCA territory at a location confirmed by a laser Doppler signal drop (**Figure 1A**). After 60 min occlusion, the filament was gently retracted, the ECA permanently ligated, the slip knot on the CCA removed, and the incision sites sutured closed. Animals were monitored for several hours post-op to ensure recovery. We have a 90% success rate of surviving animals after an effective MCAO procedure. All animals surviving to the one-day post-MCAO time point were included in the study.

**Figure 1.**
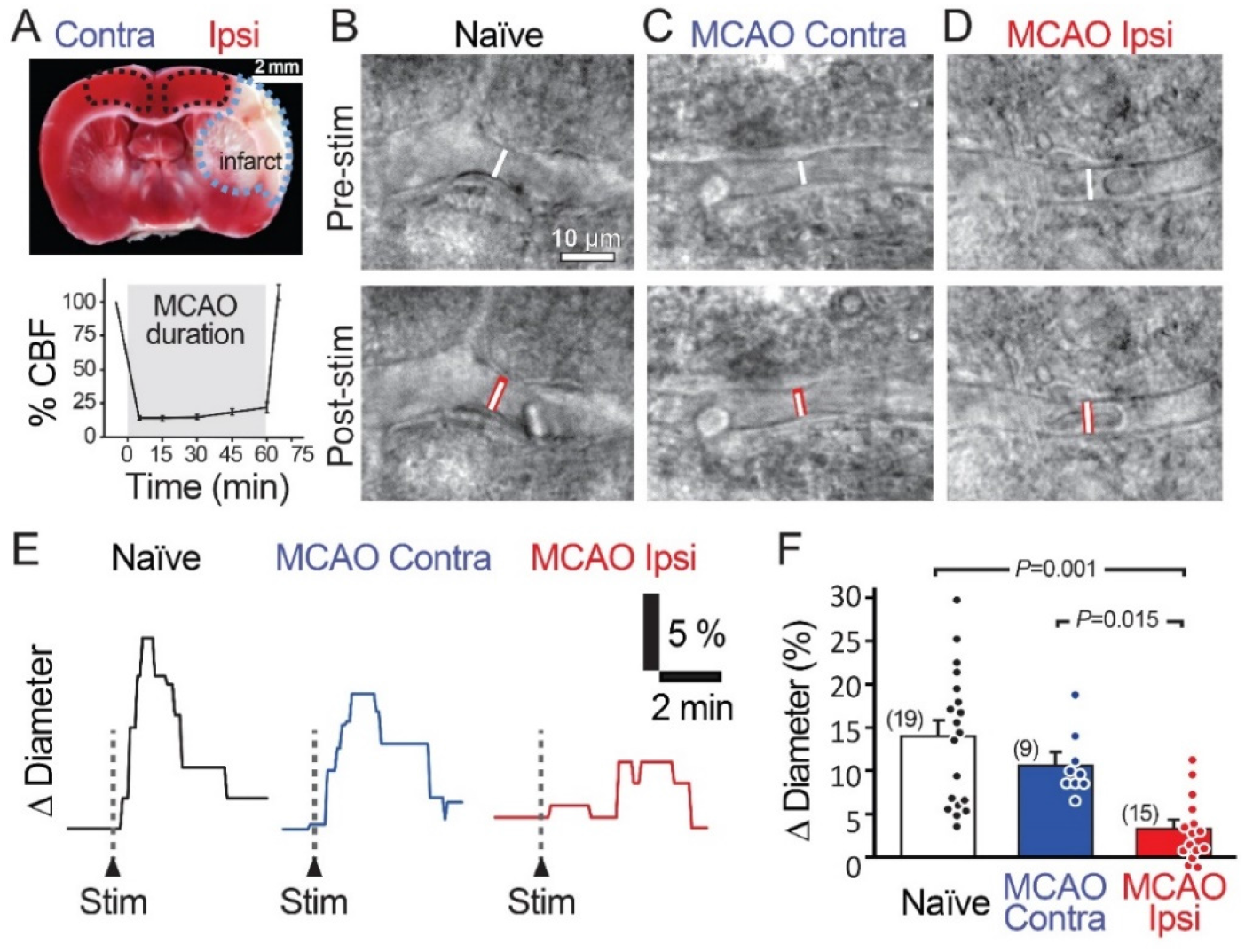
Stimulation-evoked capillary dilation is reduced in the peri-infarct cortex after MCAO. **A**. TTC staining (top) of a coronal brain slice 24 h following 60 min MCAO showing the unstained infarcted region in white (outlined in blue) and intact tissue in red. MCAO Ipsi data were acquired from the peri-infarct cortical region and contralateral hemisphere and naïve brain slice data were acquired in the analogous region (outlined in black). Bottom graph shows the decrease in cerebral blood flow (CBF) in the MCA territory during occlusion. **B-D**. An example capillary in naïve **(B)**, MCAO Contra **(C)** and MCAO Ipsi **(D)** cortex before (top) and after (bottom) neuronal stimulation. Dotted white lines show capillary diameter in U46619 prior to stimulation, while the underlaid red bars in lower panels show capillary diameter after stimulation. **E-F**. Example traces **(E)** and summary quantification **(F)** demonstrating the reduction in stimulation-evoked capillary dilation in the MCAO Ipsi peri-infarct cortex. *P* values were obtained from ANOVA followed by Tukey’s HSD test. Number in parentheses above each bar indicates *N*.

### Brain Slice Preparation

Acute brain slices were prepared as described previously [30] from rats one day after MCAO surgery. Briefly, 300-μm-thick coronal cortical slices were prepared on a Vibratome in ice-cold oxygenated (95% O_2_, 5% CO_2_) solution containing (in mM) 93 *N*-methyl-D-glucamine (NMDG) chloride, 2.5 KCl, 30 NaHCO_3_, 10 MgCl_2_, 1.2 NaH_2_PO_4_, 25 glucose, 0.5 CaCl_2_, 20 HEPES, 5 Na ascorbate, 3 Na pyruvate, 2 thiourea, and 1 kynurenic acid. The slices were incubated at 34°C in the same solution for 15–20 min, then transferred to a similar solution with the NMDG chloride, MgCl_2_, and CaCl_2_ replaced by (mM) 93 NaCl, 1 MgCl_2_, and 2 CaCl_2_, and incubated at room temperature (21°C–23°C) for at least 20 min or until used in experiments (maximum of 5 h).

### Identification and Imaging of Capillaries

Slices were perfused with bicarbonate-buffered artificial cerebrospinal fluid (aCSF) containing (in mM) 124 NaCl, 2.5 KCl, 26 NaHCO_3_, 1 MgCl_2_, 2 CaCl_2_, 1 NaH_2_PO_4_, 10 glucose, 1 Na ascorbate, warmed to 32°C-34°C and bubbled with 20% O_2_, 5% CO_2_, and 75% N_2_ to mimic physiologic O_2_ conditions and avoid O_2_ or metabolism-dependent modulation of vascular responses [22, 31, 32]. Imaging was performed using differential interference contrast (DIC) microscopy with a 40X water immersion objective, a CMOS USB 3.0 camera (Thorlabs), and Micro-Manager acquisition software. Capillaries were imaged at 15–50 μm depth in layers III–VI of slices containing the sensorimotor cortices, in the peri-infarct region (stroke hemisphere) or the analogous region (contralateral and naïve slices) (**Figure 1B**). The peri-infarct region was defined as all dorsomedial cortical regions >0.5 mm away from the infarct border, which is identified as the area comprising dead and swollen cells. Capillaries were defined, as previously described, as vessels with <8 μm luminal diameter and lacking abluminal contiguous smooth muscle cells, often instead containing pericytes identified by their bump-on-a-log morphology at ∼30 μm intervals [30]. Images were acquired every 5 s, with 10 ms exposure, at a resolution of 132 nm/pixel. As vessels in acute brain slices lack perfusion and therefore tone, U46619 (200 nM) was applied in the bath for at least 5 min to preconstrict capillaries in all experiments assaying neurovascular coupling. Pre-constriction to U46619 ensured that the vessel was healthy and responsive, therefore lack of constriction to U46619 was an exclusion criterion [20]. Vessel internal diameters were measured by manually placing a measurement line perpendicular to the vessel on the image at locations where constriction to U46619 occurred (presumed regions of active pericyte control) using MetaMorph software. Change in diameter was quantified as a 30 s average centered around the largest response seen after stimulation, normalized to baseline diameter. If changes in focus confounded accurate measurements, experiments were excluded from further analysis. Because of the obviousness of the MCAO’s effects on both animal behavior (circling in cage) and tissue health within the infarct in acute slices, experiments could not be performed blind. The need to keep track of the ipsilesional and contralesional slices in MCAO group further precluded blinding during experiments. Thus, experiments were interleaved between the stroke hemisphere and the contralateral slices from each animal to minimize confounds due to day-to-day and time-from-slicing variability. Experiments in naïve slices were performed on alternate days. To minimize bias, all experiments were conducted by one experimenter and the resulting files were analyzed by an independent investigator blinded to the conditions.

### Stimulation Protocol for Brain Slices

Electrical stimulation was applied using an aCSF-filled glass electrode with a wide opening placed in layer I/II of the slice, approximately 300–500 μm from the imaged vessel. Two separate stimulation paradigms were tested (**Supplementary Figure 1C-D**): a 1 s, 100 Hz (high-frequency), and a 3 s, 20 Hz (low-frequency) stimulation with 0.2 ms pulse duration and 200 mA intensity using a constant current stimulator (Digitimer). Both paradigms evoked capillary dilations with the same pharmacology (blocked by NF449) as previously described [20]. The low-frequency stimulation was chosen as the physiologically relevant minimal stimulation to evoke neuronal activity in all experiments (**Figures 1-4**). To confirm activation of the neurons, extracellular field recordings were performed near the capillary of interest using a 3-5 MΩ glass electrode filled with aCSF, and only experiments where an fEPSC was detected were used for analysis. The amplitude and slope of the first fEPSC were quantified to compare neuronal activation.

### Pharmacology in Brain Slices

For all slice experiments, vessels were pre-constricted with U46619, which acts on thromboxane A_2_ receptors and constricts capillaries without altering neuronal activity [20]. U46619 was applied for at least 5 min before any further manipulations (application of drugs or electrical stimulation). Capillaries that did not constrict to U46619 were disqualified from analysis to exclude vessels in which contractility may be compromised. The drug of interest was then bath applied for at least 5 min before stimulation-evoked capillary responses were evaluated. Each drug was used at a concentration range within two-to ten-fold higher than its EC_50_/IC_50_ to ensure its effects without introducing non-specificity.

### Arterial spin-labeling MRI - Data Acquisition

All magnetic resonance (MR) imaging was performed on a Bruker BioSpin 11.75 T (Bruker Scientific Instruments, Billerica, MA) small animal MR system with a horizontal bore, 9-cm inner diameter (ID) gradient set (750 mT/m), with a 72 mm ID and 60 mm (length) radiofrequency resonator and an actively decoupled 20 mm Bruker surface coil for transmit/receive. Rats were anesthetized with mask isoflurane (∼1.5%) in 100% oxygen and subjected to lateral tail vein canulation with a 27-gauge Terumo Surflo catheter (Thermo Fisher Scientific). A custom platform/head holder was used to dorsoventrally position the rats and immobilize the head. Body temperature was maintained at 37°C using a biofeedback warm air temperature control system (Small Animal Instruments, Inc.). Respiration rate was monitored using the MR-compatible Monitoring & Gating System (Small Animal Instruments, Inc.) and maintained between 75-110 breaths/min by slight tuning of isoflurane level. A coronal T_2_-weighted 35-slice image set was obtained (spin echo RARE, 256 × 256 matrix, 138×138 µm in-plane resolution, field of view (FOV) = 3.5 × 3.5 cm, 0.5 mm slice thickness, TR = 4065 ms, TE_effective_ = 23.6 ms, RARE factor 8, 2 averages). CBF was measured using ASL with flow-sensitive alternating inversion-recovery rapid acquisition with relaxation enhancement pulse sequence (FAIR-RARE), with echo time/repetition time (TE/TR) = 45.2 ms/10000 ms, FOV = 3.5 × 3.5 cm, slice thickness = 2 mm, number of slices = 1, matrix = 128 × 128, RARE factor = 72, and 23 inversion times ranging from 40 to 4400 ms. This sequence labels the inflowing blood by global inversion of the equilibrium magnetization [33] with a total acquisition time of 15 min. In rats exposed to MCAO, the T_2_-weighted images were used as a localizer to accurately place the ASL slice at the coronal position where maximal infarct volume was observed. For control (naïve) rats, coronal slices known to contain brain regions supplied by the MCA that were comparable in anatomy to those used in MCAO rats were chosen. Following first image capture, volume equivalent doses of either HET0016 (1 mg/kg) or vehicle (DMSO, approximately 1.8% in saline) were then slowly infused over 1 min via tail vein catheter. ASL imaging was repeated 30 min after injection. As the infarct in the MCAO group was obvious under T_2_-weighted images, blinding during ASL-MRI was not possible.

### Arterial Spin-labeling MRI - Image Analysis

Jim 7.0 image analysis software (Xinapse Systems, UK) was used for image processing. CBF maps (units: mL/100 g/min) were generated using the Bruker ASL perfusion processing macro, which uses the equation:

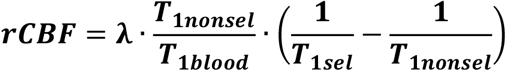

where λ = the blood-brain partition coefficient (0.9), T_1blood_ =the T_1_ of blood (2800 ms), and T_1sel_ and T_1nonsel_ are the measured T_1_ values under selective and nonselective inversion conditions, respectively. CBF maps were exported into Jim imaging software (Version 8.0) for further processing. The ROI (region of interest) toolkit was used to manually outline the entire brain on a paired T_2_-weighted image slice, which was subsequently divided into left and right hemispheres. Next, the ipsilateral peri-infarct cortex and contralateral analogous cortex ROIs were determined based on the T_2_-weighted image for each animal taking care to avoid the infarct lesion and the underlying white matter (corpus callosum). The identical FOV and geometry of the T_2_ and ASL volumes enabled the ROIs to be readily overlaid onto the corresponding perfusion map for quantification. To remove flow contributions from large vessels and better quantify capillary flow, voxels with intensities +/-2 SD of the mean were excluded. The average of all remaining voxel intensities was then used to calculate corrected flow values within an ROI. To quantify the change in CBF induced by the treatment (vehicle or HET0016), flow values for each ROI after the treatment injection were normalized to pre-treatment values.

### Preparation of Samples for 20-HETE and PGE2 Measurement

20-HETE and PGE2 are both lipid signaling molecules that are synthesized on-demand and can easily pass the membrane to signal to other cells; thus, it is most appropriate to measure whole tissue lysate concentrations. Brains from rats treated with vehicle or HET0016 during the ASL experiment were dissected within 10 min after MRI and 500-μm thick live brain slices were cut using a Vibratome (Leica) spanning the sensorimotor cortex within the MCA territory, as determined visually based on comparison with triphenyl tetrazolium chloride (TTC) stains on other MCAO brains (to define stroke region), the T_2_ images from the same animal, and the Rat Brain Atlas (Paxinos and Watson). Immediately after cutting, the stroke peri-infarct and analogous cortical regions (from control slices) were carefully microdissected from each slice, excess aCSF was gently wicked away from the tissue with a Kimwipe™, and the tissue was transferred to a vial containing 1 mL of 0.1% formic acid in water. This was performed in duplicate for each vial, such that each sample contained microdissected cortical regions from two adjacent coronal slices. Two vials were prepared from each hemisphere of each animal and immediately placed in a -80°C freezer to preserve tissue contents. The frozen vials were blinded and transferred to the Bioanalytical Shared Resource/Pharmacokinetics Core at Oregon Health & Science University.

At the time of measurement, vials were thawed and the tissue homogenized using a BeadBug microtube homogenizer. A 0.9 mL aliquot of the tissue homogenate was placed into a 13×100 mm glass screw-top extraction tube with a Teflon™-lined cap. The remaining 0.1 mL of the homogenate was used for protein quantification for normalization of the data. Deuterated internal standards were added (1 ng of PGE2-d_9_ and 2 ng 20-HETE-d_6_) to the homogenate and the tubes were extracted in sequence with 3 mL of ethyl acetate with 0.02 mg/mL triphenylphosphine, followed by 3 mL of a 1:1 mixture of ethyl acetate:hexane, and finally with 2 mL of hexane. The combined supernatants were spiked with 20 μL of a trap solution consisting of 10% glycerol in methanol with 0.01 mg/mL butylated hydroxytoluene. The samples were dried for 45 min in a speed vacuum at 35°C, the walls of the tubes were washed with 1 mL of hexane and re-dried until a small aqueous residue remained. The residue was dissolved in 80 μL of 80:20 water:acetonitrile with 0.1 mg/mL butylated hydroxytoluene and spin filtered with a 0.22-μm Millipore ultra-free filter. Samples were transferred to vials and 30 μL of sample was analyzed by LC-MS/MS as described below. Sample extraction buffer was spiked and prepared identically to the samples with concentration ranges from 10 to 2500 pg/mL for PGE2 and 20-HETE.

### Liquid Chromatography Followed by Tandem Mass Spectroscopy (LC-MS/MS)

Extracts were analyzed using a 5500 Q-TRAP hybrid/triple quadrupole linear ion trap mass spectrometer (SCIEX, Carlsbad, CA) with electrospray ionization (ESI) in negative mode. The mass spectrometer was interfaced to a Shimadzu (Columbia, MD) SIL-20AC XR auto-sampler followed by two LC-20AD XR LC pumps. The scheduled MRM transitions were monitored with a 1.5 min window. Optimal instrument parameters were determined by direct infusion of each analyte and are presented in the table below.

| Analyte | Q1 | Q3 | RT | DP | EP | CE | CXP |
| --- | --- | --- | --- | --- | --- | --- | --- |
| PGE2 | 351.0 | 189.1 | 5.9 | -85 | -10 | -26 | -5 |
| PGE2-d<sub>9</sub> | 360.2 | 280.2 | 5.9 | -75 | -10 | -28 | -11 |
| 20-HETE | 319.1 | 289 | 10.3 | -130 | -10 | -26 | -11 |
| 20-HETE-d<sub>6</sub> | 325 | 281 | 10.3 | -75 | -10 | -24 | -9 |

Q1 = Quadrupole Mass filter 1; Q3 is the Quadrupole Mass filter 3; RT = retention time; DP = declustering potential; EP = entrance potential; CE = collision energy; CXP = collision cell exit potential; PGE2-d<sub>9</sub> and 20-HETE-d<sub>6</sub> = deuterium substituted versions of each compound used as internal standards.

The gradient mobile phase was delivered at a flow rate of 0.5 mL/min and consisted of two solvents, 0.05% acetic acid in water and acetonitrile. The analytes were resolved on a BetaBasic-C18 (100×2 mm, 3 µm) column at 40°C using the Shimadzu column oven. Data was acquired using Analyst 1.5.1 and analyzed using MultiQuant 3.0.1 (AB Sciex, Ontario, Canada). The lower limit of quantification was 10 pg/mL for both PGE2 and 20-HETE. As samples were blinded before sending to the core laboratory, all measurements and analyses were performed blindly by default.

### Protein Quantification

Protein quantification was performed using the Qubit 3.0 Fluorometer (Invitrogen/ThermoFisher) on the 0.1 mL of homogenate prepared in 0.1% formic acid, as described in ‘Preparation of samples for 20-HETE and PGE2 measurement’ section. All reagents and materials were brought to room temperature before mixing to ensure optimal fluorescence and measurement accuracy. Qubit working solution and Qubit Standards 1, 2, and 3 were made fresh and calibration was performed using the fresh standards. Qubit working solution was prepared by diluting Qubit Protein Reagent 1:200 in Qubit Protein Buffer in a 15 mL Eppendorf. For each of the three standards and all samples, 190 µL of working solution and 10 µL of each standard were mixed by vortexing for 2-3 seconds. All samples rested at room temperature for 15 min following preparation. All readings were performed using the “Protein” function of the Qubit 3.0 fluorometer. Standard calibration was performed by measuring Standard 1, Standard 2, and Standard 3, in order. Only calibrations that produced a linear fluorescence curve were used for sample reading. The samples were split into multiple batches of 35 samples each, which allowed for the most accurate sample readings compared to the standards calibration. Immediately after calibration, each sample was read in order. Following completion of all samples in a single batch, samples were re-run from the beginning to determine the consistency of readings. The first and second readings were in close agreement. The average of both readings was used for all final comparisons.

### Statistical Analyses

In all figures, data are shown as mean ± s.e.m. with the raw data values indicated by individual points and the *N* indicated in parentheses. In our experience, the variability of responses observed at different regions (presumed regions of active pericyte control) along the same capillary is equal to that between different capillaries, slices, or animals. Therefore, capillary regions that constricted to 200 nM U46619 were used as the observational units in all slice experiments. *N* indicates the number of pericyte regions studies in **Figures 1-4** and **Supplemental Figure 1**, the number of animals in **Figure 5C-D**, and the number of samples in **Figure 5E-F**. Data from 3-6 rats were used for all experiments. Normality of data was checked using the Kolmogorov-Smirnov test or d’Agostino-Pearson test and the equality of variance using the *F*-statistic. All normal data were compared using an Analysis of Variance (ANOVA) followed by Tukey’s honest significant difference (HSD) test. Data in **Figure 5F** were non-normal, and therefore a Kruskal-Wallis test was used followed by Mann-Whitney test and Holm-Bonferroni multiple comparisons correction. Two-way ANOVA was used in **Figure 5C** to test whether a significant interaction between stroke exposure and hemispheric CBF could be detected. Sample sizes were calculated based on previous experience with the experimental techniques used and expected variability.

## Results

To assess neurovascular coupling in acute brain slices, we monitored changes in capillary diameter evoked by electrical stimulation of cortical neuronal activity [20]. We first confirmed that neuronal activity evoked capillary dilation in adult (2-month-old) rat cortical slices by applying two different stimulation paradigms. We found that both high-frequency (100 Hz, 1 s) stimulation and low-frequency (20 Hz, 3 s) stimulation evoked robust capillary dilation (18.9 ± 4.7%, n=7, and 14.5 ± 1.8%, n=18, for high and low-frequency stimulation, respectively; **Supplementary Figure 1**). We previously showed that activity-dependent capillary dilation is mediated by an astrocytic signaling pathway dependent on the P2×1 receptor [20]. In our current experiments, bath application of the P2×1 inhibitor NF449 (100 nM) similarly reduced capillary dilation evoked by both high (3.9 ± 2.2%, n=6, *P=*0.01) and low-frequency (1.9 ± 1.3%, n=20, *P=*0.00002) stimulation (**Figure Supplementary 1B-D**). The low-frequency stimulation was chosen as a physiologically relevant minimal stimulation paradigm for all further experiments [34].

We next examined capillary responses evoked by low-frequency stimulation in the peri-infarct cortex of animals exposed to MCAO (MCAO Ipsi) and compared them to the analogous cortical region in contralateral slices (MCAO Contra) and slices from naïve animals (**Figure 1**). Stimulation evoked a capillary dilation of 14.0 ± 1.8% (n=19) in naïve cortex and 10.7 ± 1.3% (n=9, *n*.*s*. compared to naïve) in the MCAO Contra region, but this response was reduced to 3.3 ± 1.0% (n=15) in the MCAO Ipsi region (ANOVA *P=*0.00003; Tukey’s HSD *P=*0.001 compared to naïve, *P=*0.015 compared to MCAO Contra; **Figure 1E-F**).

Stimulation-evoked capillary responses may be reduced in the peri-infarct cortex for several reasons. The most parsimonious explanation is that neuronal activity is itself reduced. We tested this possibility by comparing the amplitude and rising slope of the stimulation-evoked field excitatory post-synaptic currents (fEPSCs; **Figure 2A,B**). The amplitude of the fEPSCs was 120.6 ± 19.6 nA (n=30) in naïve brains, 127.8 ± 18.8 nA (n=24) in the MCAO Contra region, and 100.8 ± 14.5 nA (n=30) in the MCAO Ipsi peri-infarct region (ANOVA *P=*0.6). The rising slope of the fEPSCs was 80.7 ± 18.4 nA/ms in naïve brains, 101.5 ± 17.7 nA/ms in the MCAO Contra region, and 81.0 ± 11.2 nA/ms in the MCAO Ipsi peri-infarct region (ANOVA *P=*0.5). A lack of vessel response could also be due to loss of vascular contractility. As dilation is in essence an inhibition of contraction, which is the active component of vascular regulation, we assessed vascular contractility by evaluating capillary constriction evoked by the thromboxane analog U46619 (**Figure 2C,D**). We found no difference between groups: U46619 (200 nM) evoked a constriction of 12.1 ± 1.1% (n=45) in naïve brains, 12.1 ± 1.4% (n=38) in the MCAO Contra region, and 12.6 ± 1.1% (n=45) in the MCAO Ipsi peri-infarct region (ANOVA *P=*0.9). We further tested a second, stronger vasoconstrictor, endothelin 1 (ET-1, 10 nM), which also had a comparable effect across conditions (**Figure 2E,F**). ET-1 constricted capillaries by 26.7 ± 6.4% (n=15) in naïve brains, 30.1 ± 8.5% (n=8) in the MCAO Contra region, and 28.2 ± 9.4% (n=12) in the MCAO Ipsi region (ANOVA *P=*0.7). Thus, the reduction in capillary dilation in the MCAO Ipsi peri-infarct cortex was not explained by a reduction in stimulation-evoked neuronal activity or loss of vasocontractility

**Figure 2.**
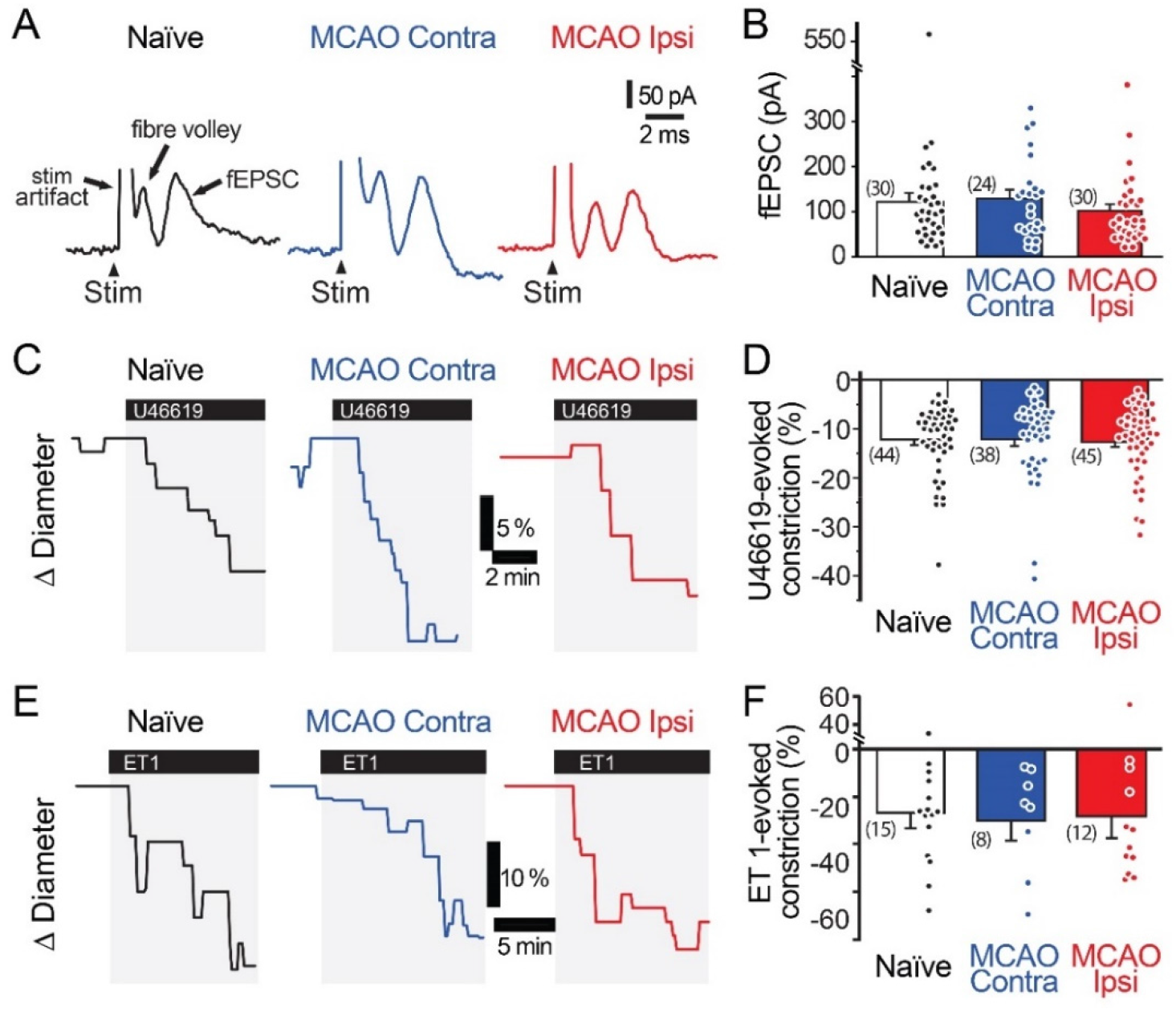
Neuronal activity and vasocontractility are not reduced in the MCAO peri-infarct cortex. **A-B**. Stimulation-evoked field excitatory post-synaptic currents (fEPSC) are comparable in the MCAO Ipsi peri-infarct and the analogous region of the MCAO Contra and naïve slices. **C-D**. 200 nM U46619 evokes a similar capillary constriction in all three regions. **E-F**. ET-1 (10 nM), a more potent vasoconstrictor, evokes a stronger but statistically similar capillary constriction in all three regions. None of the comparisons were statistically significant based on ANOVA followed by Tukey’s HSD test. Number in parentheses above each bar indicates *N*.

Increased levels of the vasoconstrictor 20-HETE in the plasma and cerebrospinal fluid correlate with worse prognosis in stroke patients [23-25]. Thus, we tested whether 20-HETE underlies the reduction in evoked capillary dilation after MCAO by incubating acute brain slices in N-Hydroxy-N′-(4-butyl-2-methylphenyl) formamidine (HET0016, 200 nM) to inhibit 20-HETE synthesis prior to assaying neurovascular coupling [35]. Pre-incubation of slices with HET0016 completely rescued capillary neurovascular coupling in the MCAO peri-infarct region (**Figure 3A,B**). Stimulation-evoked capillary dilation in the presence of HET0016 was 10.4 ± 2.0% ± 8.5% (n=8) in the (n=12) in naïve brain slices, 10.1 ± 1.8% (n=14) in the MCAO Contra region, and 11.1 ± 2.4% (n=16) in the MCAO Ipsi peri-infarct region (ANOVA *P=*0.9). HET0016 treatment did not directly alter the diameter of the vessels (**Figure 4A**).

**Figure 3.**
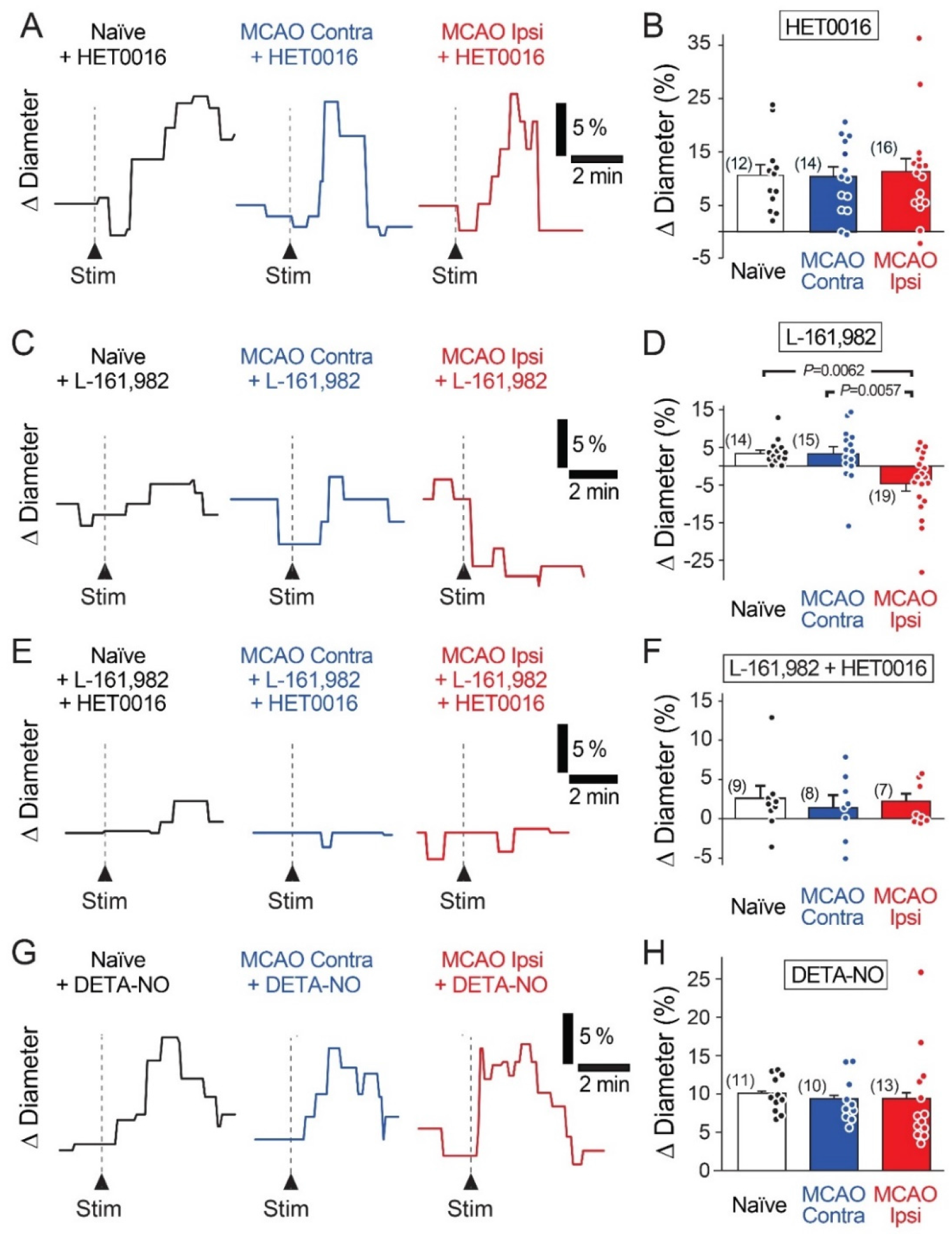
Increased 20-HETE synthesis contributes to the reduction in neurovascular coupling in the MCAO Ipsi peri-infarct cortex. **A-B**. HET0016 (200 nM), a potent inhibitor of 20-HETE synthesis, rescues neuronally evoked capillary dilations without altering responses in contralateral slices or naïve brains. **C-D**. Blocking the EP4 receptor of PGE2 with L-161,982 (1 μM) unmasks a 20-HETE-mediated constriction only in the MCAO ipsilateral peri-infarct region. **E-F**. Inhibiting EP4 receptors and 20-HETE synthesis simultaneously prevents capillary response to neuronal activation in all three conditions. **G-H**. Increasing NO concentration in the tissue with DETA-NONOate (DETA-NO, 100 μM) restores stimulation-evoked capillary dilation in the MCAO ipsilateral hemisphere without altering capillary responses in contralateral or naïve brain slices. *P* values were obtained from ANOVA followed by Tukey’s HSD test. Number in parentheses above each bar indicates *N*.

**Figure 4.**
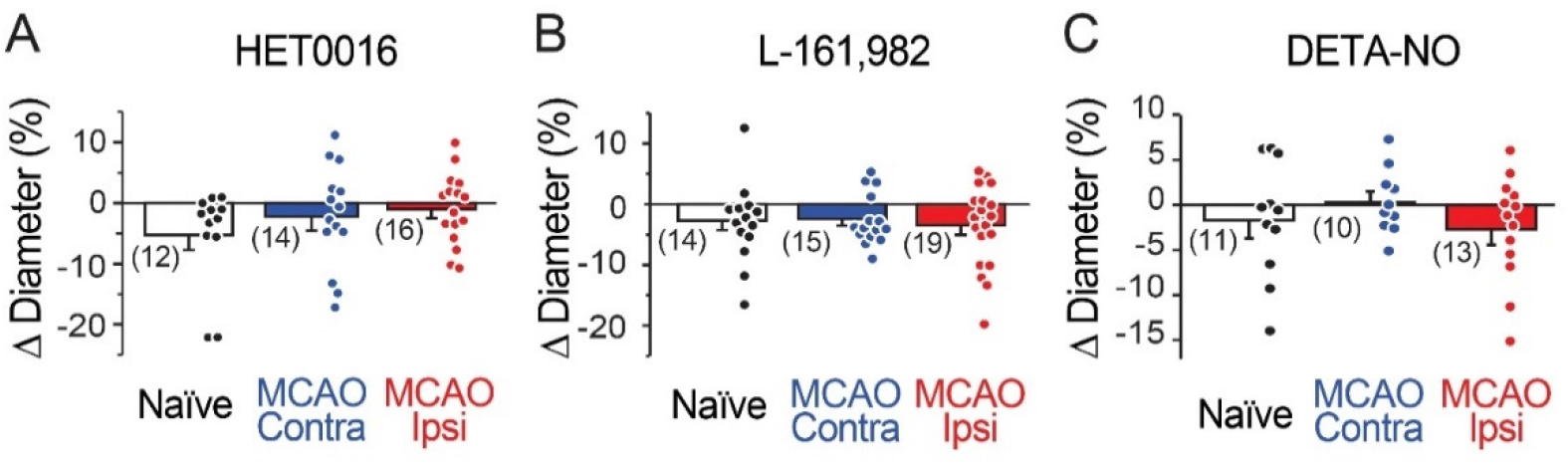
Capillary diameter is not directly altered by 20-HETE, EP4, or NO. Reducing 20-HETE synthesis with HET0016 **(A)**, blocking the PGE2 receptor EP4 with L-161,982 **(B)** or increasing ambient NO level with DETA-NONOate (DETA-NO; **C**) does not alter the resting diameter of capillaries (under U46619-mediated preconstriction).

As inhibition of 20-HETE synthesis revealed the presence of stimulation-evoked capillary dilation in the MCAO Ipsi peri-infarct cortex, we next sought to identify this dilatory signal. In healthy cortical tissue, capillary neurovascular coupling depends on PGE2 acting on its receptor EP4 [20, 22]. If PGE2-dependent vasodilatory signaling persists after MCAO but is masked by coincidental synthesis of 20-HETE, then inhibiting the PGE2 pathway should reveal a stimulation-evoked constriction in the MCAO Ipsi peri-infarct cortex due to 20-HETE. Application of the EP4 receptor blocker L-161,982 (1 µM) to acute slices reduced neuronal stimulation-evoked capillary dilation in naïve brain slices (3.3 ± 0.9%; n=14) and in the MCAO Contra region (3.2 ± 1.8%; n=15; **Figure 3C,D**). In the MCAO Ipsi region, L-161,982 treatment converted stimulation-evoked capillary responses to constrictions as predicted (−4.6 ± 1.9%; n=19; ANOVA *P=*0.002; Tukey’s HSD *P=*0.006 compared to naïve, *P=*0.005 compared to MCAO Contra). L-161,982 did not affect baseline diameter (**Figure 4B**). When both 20-HETE and PGE2 pathways were simultaneously inhibited (200 nM HET0016 plus 1 µM L-161,982), stimulation-evoked responses of either polarity were absent in all conditions (**Figure 3E,F**). The evoked change in capillary diameter measured 2.6 ± 1.5% (n=9) in naïve slices, 1.4 ± 1.5% (n=8) in the MCAO Contra region, and 2.1 ± 0.9% (n=8) in the MCAO Ipsi region (ANOVA *P=*0.8).

In healthy brains, CYP450 enzymes are inhibited by NO, which results in low levels of tissue 20-HETE [22]. A reduction in neuronal and endothelial isoforms of NO synthase, leading to a decrease in activity-dependent NO production, is reported after ischemic injury [28], which could lead to increased CYP450 activity. We reasoned that if disinhibition of CYP450s due to a decrease in NO is responsible for enhancing 20-HETE synthesis after MCAO, we should be able to rescue neurovascular coupling simply by exogenously raising NO concentration. To this end, we performed experiments in the presence of diethylenetriamine NONOate (DETA-NONO; 100 µM), a long half-life NO donor that releases a reasonably constant amount of NO over several hours [36]. When NO levels were thus raised in slices, capillary neurovascular coupling was rescued (**Figure 3G,H**) comparably to when 20-HETE synthesis was inhibited, while having no direct effect on baseline capillary diameter (**Figure 4C**). Stimulation-evoked capillary dilation in the presence of DETA-NONO was 10.1 ± 0.7% (n=11) in naïve slices, 9.4 ± 0.9% (n=10) in the MCAO Contra region, and 9.4 ± 1.7% (n=13) in the MCAO Ipsi peri-infarct region (ANOVA *P=*0.9).

We next sought to examine whether 20-HETE contributes to changes in CBF after stroke *in vivo*. We used ASL-MRI to map cortical perfusion in anesthetized rats before and 30 min after intravenous (i.v.) injection of HET0016 (1 mg/kg) or vehicle. The maps were corrected to largely reflect capillary blood flow (see Methods) and CBF was quantified in the MCAO Ipsi peri-infarct cortex and analogous regions of the MCAO Contra and naïve cortices (outlined in **Figure 5A,B**). CBF was significantly higher in the cortex of naïve rats compared to those exposed to MCAO in both hemispheres (**Figure 5C**; naïve: 36.4 ± 1.8 mL/100 g/min in Ipsi and 34.1 ± 2.0 mL/100 g/min in Contra, n=11; MCAO: 27.2 ± 2.5 mL/100 g/min in Ipsi and 27.3 ± 1.8 mL/100 g/min in Contra, n=10; two-way ANOVA *P=*0.0004 between naïve vs MCAO).

**Figure 5.**
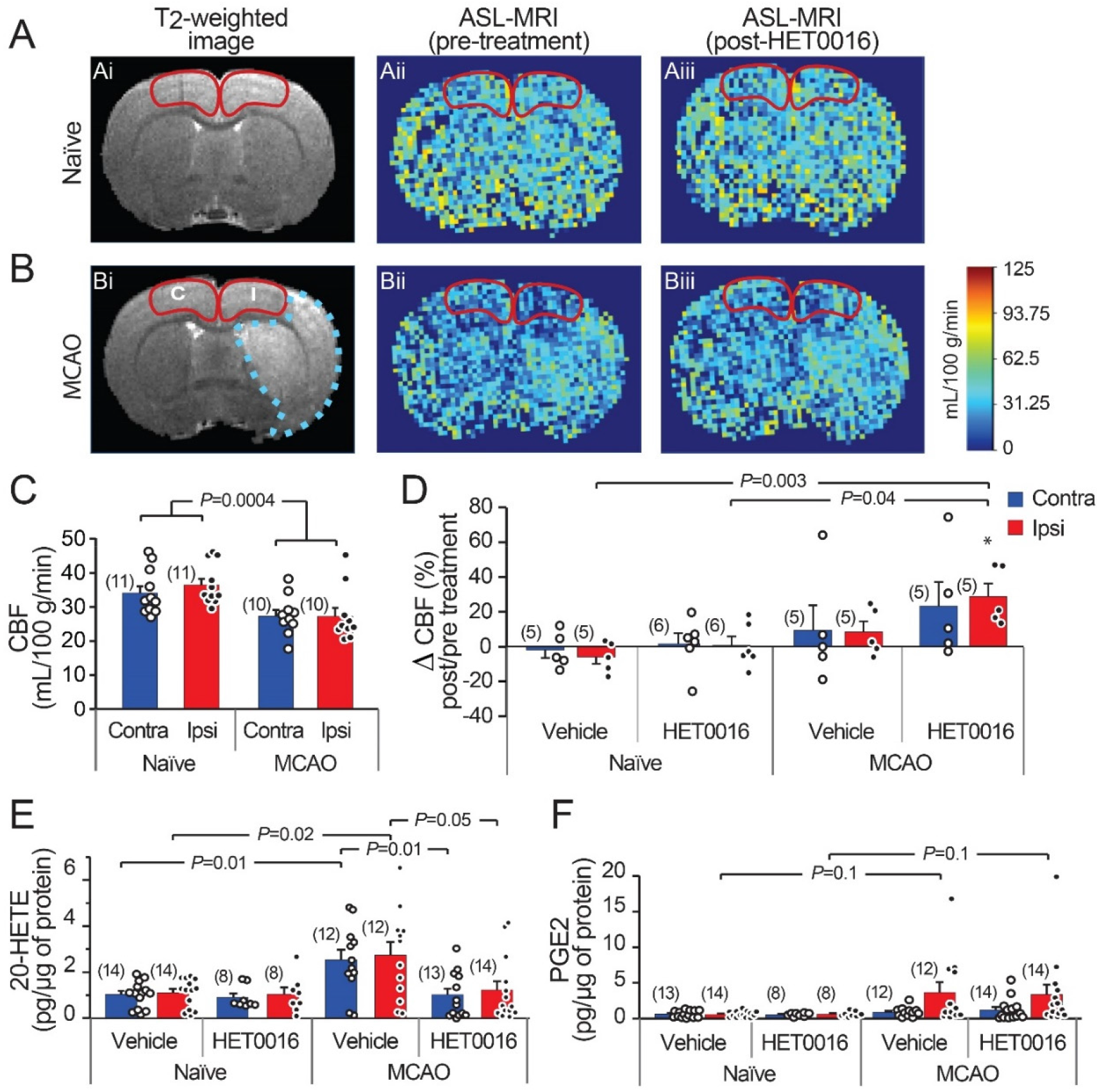
Increased 20-HETE synthesis reduces resting CBF in the stroke peri-infarct. **A-B**. T_2_-weighted MRI and ASL images of CBF in naïve and MCAO rats. CBF was regionally quantified in the cortical regions outlined in red (I = ipsilateral peri-infarct, C = contralateral). **C**. Cortical CBF is reduced bilaterally in MCAO animals compared to naïve animals. **D**. HET0016 (1 mg/g i.v.) increases CBF significantly in the MCAO Ipsi peri-infarct region without altering CBF in naïve brains. **E-F**. Mass spectrometric measurement of 20-HETE and PGE2 in microdissected cortical tissue (region marked in **A-B**), normalized to total protein. **E**. 20-HETE is increased in both contralateral and ipsilateral cortices of MCAO rats and treatment with HET0016 reduces it to baseline levels comparable to naïve rats. **F**. Cortical PGE2 levels are not changed significantly after MCAO and not altered by HET0016 treatment. *P* value in **C** obtained from a two-way ANOVA. *P* value in **D-F** obtained from ANOVA followed by Tukey’s HSD. Number in parentheses above each bar indicates *N*.

Acute delivery of HET0016 (i.v.) increased CBF in the MCAO brains bilaterally (28.8 ± 7.4% increase in Ipsi and 23.2 ± 14.1% increase in Contra), but this change reached significance only in the MCAO Ipsi peri-infarct cortex (one-sample t-test *P=*0.009 compared to zero change). The effect of HET0016 treatment on CBF in MCAO Ipsi cortex was in stark contrast to that observed in naïve cortices, where CBF changes were essentially absent (**Figure 5D**; ANOVA *P=*0.0036). These data strongly support the idea that 20-HETE contributes to a decrease in cortical blood flow following MCAO, especially in the peri-infarct region. Vehicle treatment did not alter CBF in either hemisphere in naïve or MCAO rats. The changes in CBF observed after vehicle and HET0016 treatment for all regions are reported in **Table 1**.

**Table 1.**
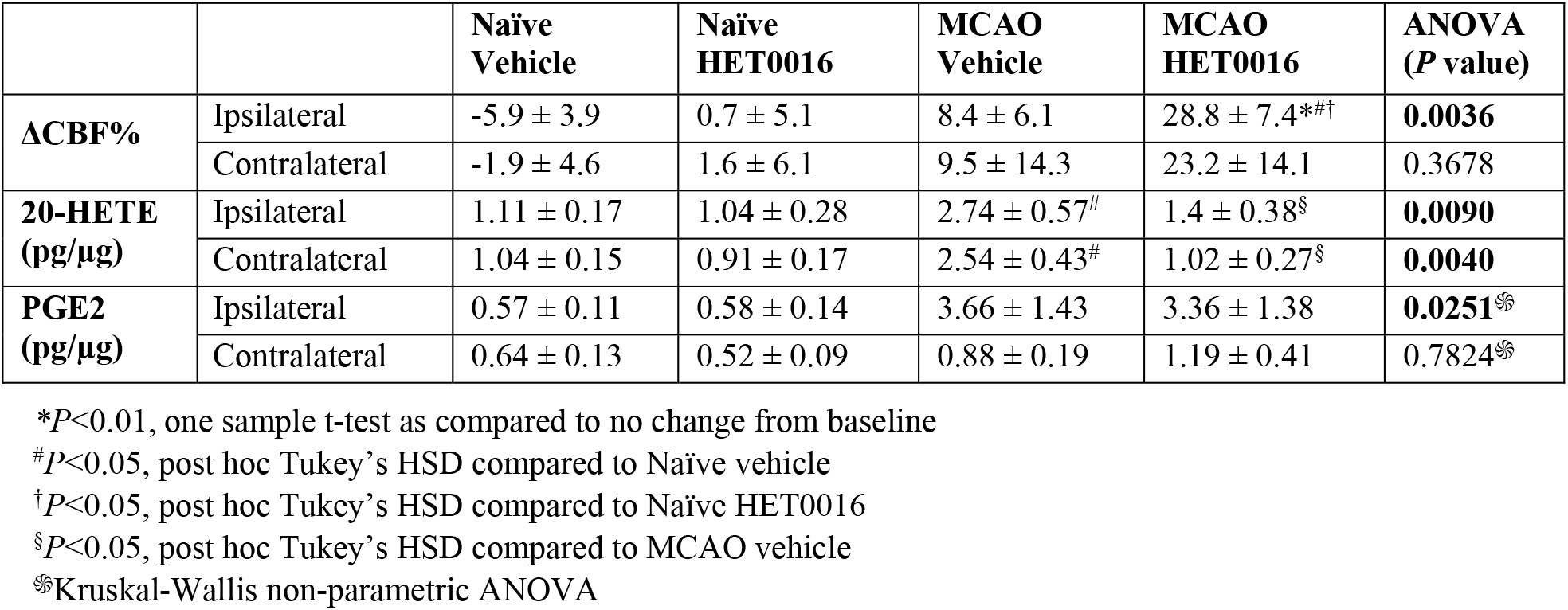
The percent change in cortical CBF measured from ASL-MRI after vehicle or HET0016 treatment compared to pre-treatment baseline (ΔCBF%). The levels of cortical 20-HETE and PGE2 measured by LC-MS/MS after vehicle or HET0016 treatment are reported as pg/µg total protein (tissue taken from the same animals in which ASL-MRI was performed, immediately following MRI). All measurements were made from the cortical region demarcated as peri-infarct (MCAO Ipsilateral hemisphere) or analogous region (contralateral hemisphere and naïve brains).

|  |  | **Naïve Vehicle** | **Naïve HET0016** | **MCAO Vehicle** | **MCAO HET0016** | **ANOVA (P value)** |
| --- | --- | --- | --- | --- | --- | --- |
| **ΔCBF%** | Ipsilateral | -5.9 ± 3.9 | 0.7 ± 5.1 | 8.4 ± 6.1 | 28.8 ± 7.4<sup>*#†</sup> | **0.0036** |
| | Contralateral | -1.9 ± 4.6 | 1.6 ± 6.1 | 9.5 ± 14.3 | 23.2 ± 14.1 | 0.3678 |
| **20-HETE (pg/μg)** | Ipsilateral | 1.11 ± 0.17 | 1.04 ± 0.28 | 2.74 ± 0.57<sup>#</sup> | 1.4 ± 0.38<sup>§</sup> | **0.0090** |
| | Contralateral | 1.04 ± 0.15 | 0.91 ± 0.17 | 2.54 ± 0.43<sup>#</sup> | 1.02 ± 0.27<sup>§</sup> | **0.0040** |
| **PGE2 (pg/μg)** | Ipsilateral | 0.57 ± 0.11 | 0.58 ± 0.14 | 3.66 ± 1.43 | 3.36 ± 1.38 | **0.0251**<sup>§§</sup> |
| | Contralateral | 0.64 ± 0.13 | 0.52 ± 0.09 | 0.88 ± 0.19 | 1.19 ± 0.41 | 0.7824<sup>§§</sup> |

<sup>\*</sup> $P < 0.01$ , one sample t-test as compared to no change from baseline
<sup>#</sup> $P < 0.05$ , post hoc Tukey's HSD compared to Naïve vehicle
<sup>†</sup> $P < 0.05$ , post hoc Tukey's HSD compared to Naïve HET0016
<sup>§</sup> $P < 0.05$ , post hoc Tukey's HSD compared to MCAO vehicle
<sup>§§</sup>Kruskal-Wallis non-parametric ANOVA

Lastly, we harvested brains from vehicle- and HET0016-treated animals immediately after MRI to measure the levels of cortical 20-HETE and PGE2 in the same cortical region where CBF was quantified, reflecting the MCAO Ipsi peri-infarct region and analogous control cortices (outlined in **Figure 5A,B**). These regions were microdissected from coronal brain slices and processed for liquid chromatography followed by tandem mass spectrometry. The values obtained for each metabolite were normalized to total protein content in each sample and are reported in **Table 1**. 20-HETE was elevated by ∼2.5-fold bilaterally in the cortex of vehicle-treated MCAO rats and acute treatment with HET0016 reduced 20-HETE to baseline levels (**Figure 5E**; ANOVA *P=*0.009 for Ipsi, *P=*0.004 for Contra). Interestingly, PGE2 levels were also increased, particularly in some samples, from the MCAO Ipsi region (non-parametric Kruskal-Wallis test, *P=*0.025), though pairwise comparison of the data did not reveal a statistical significance between any groups (**Figure 5F**). Furthermore, cortical PGE2 levels were not altered by HET0016 treatment.

## Discussion

Our experiments show that capillary neurovascular coupling is reduced in cortical regions beyond the infarct after MCAO and that this reduction is due to increased synthesis of 20-HETE, a potent vasoconstrictor. PGE2-mediated dilation of capillaries was intact after transient MCAO and inhibition of 20-HETE synthesis with HET0016 allowed healthy activity-coupled PGE2-mediated capillary dilations to occur. Increasing NO, an inhibitor of 20-HETE synthesis, can similarly rescue capillary dilation. *In vivo*, 20-HETE levels were increased by 2.5-fold in the peri-infarct cortex of MCAO brains and correlated with a decrease in cortical CBF. Inhibiting 20-HETE synthesis normalized 20-HETE levels and significantly increased CBF in the peri-infarct cortex. Our findings suggest that engagement of a 20-HETE-dependent vasoconstrictor pathway causes the loss of neurovascular coupling, at least in part, in otherwise healthy peri-infarct cortical tissue following ischemic stroke (**Figure 6**).

**Figure 6.**
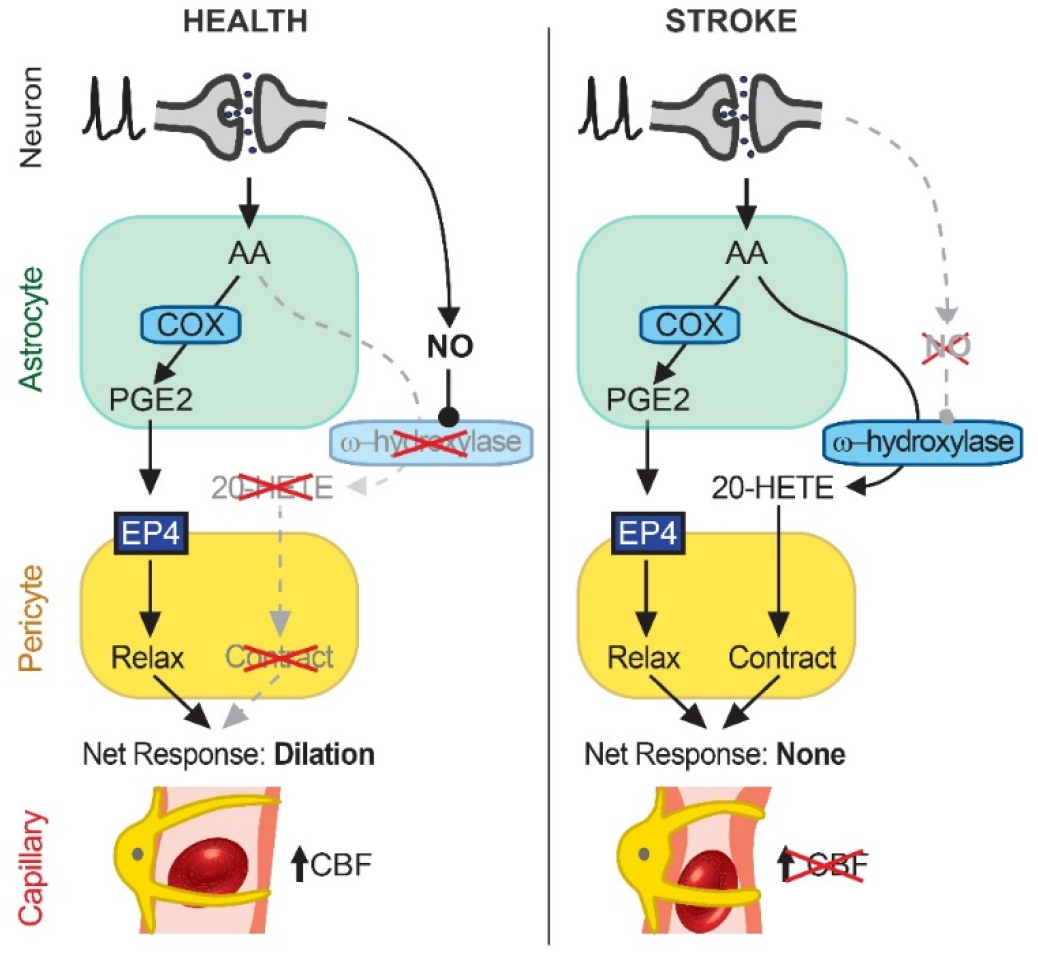
Schematic summary of findings. In healthy cortical tissue (left), neuronal activity stimulates AA synthesis and downstream metabolism to PGE2 in astrocytes. PGE2 acts on EP4 receptors on pericytes to relax them and induce capillary dilation, thus allowing an activity-coupled increase in CBF to occur. 20-HETE synthesis is inhibited by NO, likely synthesized in an activity-dependent manner from neurons (26), which suppresses pericyte contraction. After stroke (right), AA in astrocytes continues to be metabolized to PGE2, which acts on EP4 receptors to relax pericytes, but the reduction in NO generation disinhibits a parallel AA metabolism pathway via CYP450 ω-hydroxylases to produce 20-HETE, which causes pericyte contraction. The cell type(s) in which CYP450 ω-hydroxylases are active remain unknown but may include both astrocytes and pericytes. The opposing effects of EP4-dependent relaxation and 20-HETE-induced contraction of pericytes result in no net change in capillary diameter and therefore no increase in CBF. Over time, this reduced neurovascular coupling could chronically impair resting CBF and contribute to cognitive decline.

Several clinical studies have demonstrated that stroke results not only in an acute loss of blood flow, and consequently tissue, within the infarcted region but also causes widespread dysfunction of neurovascular physiology beyond the lesion [7-10]. Cerebral autoregulation, vascular reactivity, and neurovascular coupling are significantly impaired even in regions far from the infarct. Our observations suggest that an increase in 20-HETE synthesis may contribute to the observed impairment in neurovascular coupling (**Figure 3**). Previous reports showing an increase in 20-HETE in the plasma of stroke patients and in the cerebrospinal fluid of subarachnoid hemorrhage patients who develop delayed cerebral ischemia support this notion [23-25]. Pretreatment with 20-HETE inhibitors has been shown to attenuate the acute decrease in CBF (up to 240 min) following MCAO [26, 37]. Our studies imply that 20-HETE-mediated suppression of neurovascular coupling is likely contributing to this decrease in CBF and that 20-HETE’s effects on neurovascular coupling and CBF continue at least up to a day after stroke. Whether 20-HETE also contributes to autoregulation and vascular reactivity deficits, and how long these effects last in the post-stroke brain, remain unknown. Importantly however, the continued presence of a dilatory signaling pathway via EP4 suggests that inhibiting 20-HETE-dependent constriction may help restore peri-infarct neurovascular coupling without tonic effects on capillaries, which may prove a powerful and therapeutically relevant tool for future drug development efforts.

Several potential mechanisms could explain the observed increase in 20-HETE synthesis. The two foremost mechanisms are an increase in CYP450 ω-hydroxylase activity or upregulation of CYP450 ω-hydroxylase enzymes after stroke. In our *ex vivo* experiments, we observed that inhibiting 20-HETE synthesis had no effect on baseline capillary diameter (**Figure 4**), despite significantly enhancing stimulation-evoked dilation in the peri-infarct region (**Figure 3A,B**). This suggests that 20-HETE synthesis occurs in an activity-dependent manner. Furthermore, increasing NO levels to enhance inhibition of CYP450 ω-hydroxylases also rescued neurovascular coupling to its full extent (**Figure 3G,H**). These data support the former hypothesis—that the increase in 20-HETE synthesis is due to increased enzymatic activity—but do not entirely exclude the latter. Several CYP450 enzymes can ω-hydroxylate arachidonic acid to generate 20-HETE [38]. Although the identity of these enzymes remains unknown, CYP4A and CYP4F have been implicated in 20-HETE synthesis in the brain [38]. However, without selective reagents to study these enzymes (e.g., antibodies, inhibitors), determining the specific ω-hydroxylase(s) involved remains challenging. Future unbiased research using transcriptomic and proteomic analysis will be necessary to elucidate which ω-hydroxylase enzymes are expressed in the brain and possibly altered after stroke. Additionally, an oversupply of the substrate due to increased production of arachidonic acid could also increase 20-HETE production.

The cellular source of 20-HETE is also not entirely clear. Astrocytes are postulated to be the major source of arachidonic acid in the brain [39] and they express the necessary enzymes [20]. A recent study showed that the 20-HETE synthesizing enzyme CYP4A is expressed in astrocytes, vascular smooth muscle cells (VSMCs), pericytes, and endothelial cells, while the recently identified 20-HETE receptor GRP75 is present on VSMCs and pericytes [40]. Being lipid-soluble, arachidonic acid could easily diffuse out of astrocytes and into other nearby cells for further metabolism. Expression of CYP4A isoforms increases in both astrocytes and VSMCs after ischemia [41, 42]. Thus, 20-HETE synthesis in astrocytes, VSMCs, and pericytes may all potentially contribute to neurovascular coupling deficits. A recent human vascular transcriptome analysis showed that several other CYP450 enzymes capable of synthesizing 20-HETE are also present in cells within the neurovascular unit [43]. In particular, CYP4Z1 is expressed in human pericytes while CYP4F3 and CYP4F11 are expressed in hippocampal and cortical astrocytes. Interestingly, CYP4Z1 (in pericytes) and CYP4F3 (in astrocytes) were both increased over two-fold in AD patients [43]. Whether this results in an increased 20-HETE burden and contributes to the vascular dysfunction observed in AD is an avenue for future research.

The fact that CYP4A expression increases in astrocytes and VSMCs after ischemia and that 20-HETE appears necessary for mounting an appropriate angiogenic response [41] suggests that 20-HETE may be beneficial in some contexts after stroke. However, prolonged overproduction of 20-HETE and its negative effects on neurovascular coupling could render this response detrimental. A better understanding of the balance between the beneficial and harmful effects of 20-HETE may help determine when and to what extent its synthesis should be inhibited post-stroke to maximize clinical outcomes.

Our findings show that neurovascular coupling is reduced only in the ipsilesional peri-infarct regions of rats exposed to MCAO, one day after the ischemic insult. Yet, we found that 20-HETE increases in both hemispheres (**Figure 5E**) and contributes to lower CBF bilaterally after MCAO (**Figure 5C-D**). 20-HETE has important roles in many other signaling processes, including a dual role in the initiation and resolution of inflammation [44]. The presence of 20-HETE in the contralateral hemisphere at the early time point studied herein may be related to an inflammatory response rather than vascular regulation. Further, we used whole tissue homogenates to measure 20-HETE levels with the assumption that tissue amounts should reflect the overall biological availability of an easily diffusible lipid signaling mediator. However, the cellular location of 20-HETE synthesis and action could determine which specific physiological processes it can alter. Notably, impairments in neurovascular coupling in stroke patients have also been reported either exclusively ipsilateral to the lesion [8], or bilaterally, both in the lesioned and unlesioned hemispheres [9, 10]. The reduction in neurovascular coupling may initially only occur in the stroke hemisphere but then spread to the contralateral hemisphere at later time points. Our finding that 20-HETE is increased bilaterally may predict a worsening of neurovascular coupling over time, in particular spreading to the contralateral hemisphere, and future work is needed to test this suggestion. Changes in neurovascular coupling at this early period after stroke could also be occurring at the level of larger vessels, which were not tested in our study but may explain the difference between our *ex vivo* observations in neurovascular coupling (capillary focused) and our *in vivo* observations in global CBF (which was optimized to largely, but not exclusively, reflect capillary flow). Although our studies focused on 20-HETE-dependent suppression of neurovascular coupling, 20-HETE may not be the sole driver of this impairment; thus, other mechanisms should also be investigated in future studies.

The absolute values of CBF measurements obtained from ASL-MRI depend on the specific ASL approach and radiofrequency coil technology used, and therefore vary between studies conducted in humans and rodents. The flow-sensitive alternating inversion-recovery (FAIR) ASL approach uses a global inversion pulse to label incoming blood with an efficiency that depends on the radiofrequency transmitter coil coverage caudal to the brain [45]. The transmitter coil length used in this study (6 cm) generally results in near-complete inversion for mouse brains but only partial inversion for rat brains, thus producing lower CBF measurements in rats [46]. The absolute values of CBF obtained in our study fell within the range reported in other rat studies employing comparable radiofrequency coil technology [47]. Changes in animal placement under the coil can also impact the values obtained; however, we took extreme care to center the head consistently within the volume transmitter coil to optimize inversion labeling and enable robust comparison between animals. Additionally, the strength of our analysis lies in the fact that the effects of HET0016 (and vehicle) were quantified as a change in CBF within the same region in each animal before and after treatment to preclude any confounds introduced by the imaging technology. Our perfusion quantification methods were further optimized to reflect microvascular flow, where we see a significant decrease after MCAO. This agrees with the reduced capillary perfusion observed even after recanalization in stroke patients [48].

In conclusion, our findings suggest that ischemic stroke suppresses activity-dependent capillary dilation in peri-infarct regions beyond the primary infarct, possibly providing a mechanistic basis for the long-lasting cerebrovascular deficits and neurological deterioration observed in stroke patients. This reduction in neurovascular coupling appears to be the result of an increase in 20-HETE synthesis by ω-hydroxylases, which masked an otherwise healthy vasoactive response governed by PGE2 signaling. The persistence of the dilatory signaling pathway further suggests that it may be possible to pharmacologically restore neurovascular coupling following stroke.

## Acknowledgments

We thank Dr. David Attwell, Dr. Steve Sullivan, Dr. Ozama Ismail, and Mr. Evan Calkins for helpful comments on the manuscript. This work was supported by a Collins Medical Trust grant and National Institutes of Health (NIH) NINDS grant (R01NS110690) to A.M.; a Western Light Talent Training Fellowship to Z.L.; an NHLBI Ruth L. Kirschstein National Research Service Award T32 (T32HL094294) to H.M.; and a Lacroute fellowship to T.L.S. The ASL-MRI experiments were performed in the Advanced Imaging Research Center Core Facility and the LC-MS/MS analysis was conducted in the Bioanalytical Shared Resource/Pharmacokinetics Core Facility, both of which are supported in part by the University Shared Resource Program at Oregon Health & Sciences University.

## Supplemental Materials

**Supplementary Figure 1.**
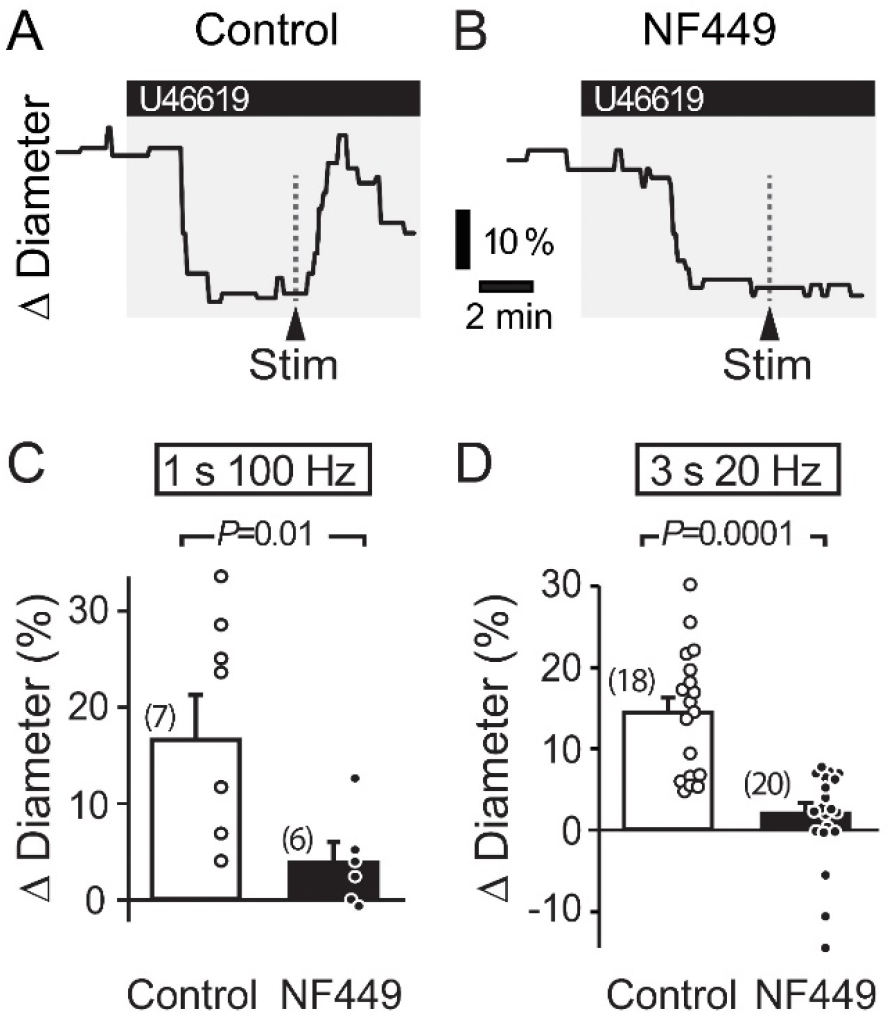
Stimulation-evoked capillary dilation in adult rat cortex depends on P2×1 receptors. **A-B**. Capillary diameter changes in response to 200 nM U46619 and superimposed neuronal stimulation (Stim; 1s 100 Hz) in a control condition **(A)** and in the presence of 100 nM NF449, a potent P2×1 blocker **(B). C-D**. Summary data demonstrating the effect of NF449 on capillary diameter evoked by a 1 s 100 Hz high-frequency stimulation (**C**) and a 3 s 20 Hz low-frequency stimulation **(D)**. *P* values obtained from Student’s t-tests. Number in parentheses above each bar indicates *N*.

